# FOSL2 Gene Expression Declines with Age in Bone Marrow-derived MSCs

**DOI:** 10.1101/2024.05.11.593704

**Authors:** James Utley, Daniel Briggs

**Affiliations:** Auragens

**Keywords:** FOSL2, Aging, Gene Expression, AP-1 Transcription Factor, Linear Regression Analysis

## Abstract

The FOSL2 gene, integral to the AP-1 transcription factor complex, orchestrates cellular responses to stimuli, including immune surveillance and tissue-resident memory T cell differentiation. This study investigates FOSL2’s expression dynamics across ages to elucidate its role in aging and age-associated diseases. Leveraging gene expression profiles in response to environmental challenges, we hypothesize that FOSL2 serves as a critical regulator of aging-associated cellular alterations. Utilizing quantitative PCR and RNA sequencing, we charted FOSL2 expression in human bone marrow-derived mesenchymal stromal cells (hMSCs) aged 17-84 years. Statistical analyses reveal a significant negative correlation between FOSL2 expression and age (slope: -0.02442, R-value: -0.41759, P-value: 0.00081), suggesting FOSL2 as a potential biomarker for aging and its involvement in the decline of regenerative capacity. The observed decrease in FOSL2 expression aligns with its role in regulating cellular processes critical in aging. Understanding FOSL2’s regulatory network offers insights into aging mechanisms and therapeutic targets for age-related diseases.

## Introduction

The FOSL2 gene, an integral component of the AP-1 transcription factor complex, is pivotal in orchestrating cellular responses to diverse external stimuli, including pathogens and ionizing radiation. Recent studies underscore its significance in the differentiation and priming of tissue-resident memory T cells (TRM), highlighting FOSL2’s role in immune surveillance and rapid reactivation of these cells in response to antigens (Smith et al., 2023). Furthermore, FOSL2, as part of the AP-1 complex, has been implicated in the transcriptional activation of senescence genes and the regulation of chromatin accessibility, which are critical processes in cellular aging (Ferreira et al., 2023). This complex interplay suggests that FOSL2’s expression patterns across different ages may offer insights into its contributions to the aging process and the development of age-associated diseases. Notably, the transcriptional activation of AP-1 complex members, including FOSL2, emerges as a conserved signature of immune aging, potentially contributing to increased inflammation and senescence, a phenomenon often referred to as “inflammaging” (Karakaslar et al., 2022). This study aims to examine the expression dynamics of FOSL2 across various life stages to elucidate its implications in aging and related pathologies, leveraging insights from gene expression profiles in response to environmental challenges (Nishad & Ghosh, 2016) and the conserved transcriptional mechanisms underlying immune cell aging (Karakaslar et al., 2022).

Hypothesis: Building on the premise that hMSCs exhibit discernible signs of aging, we postulate that the FOSL2 gene serves as a critical regulatory element in aging-associated cellular alterations. This gene’s expression profile may not only provide a molecular signature of aging but also offer a strategic target to rejuvenate hMSCs or select more efficacious cells for therapeutic applications, thus addressing the challenges posed by the inherent variability in donor-derived cells’ regenerative potential.

## Objective

To investigate the changes in FOSL2 gene expression across aging and to identify its potential impact on cellular processes and age-related diseases.

Gene Expression Dynamics: Utilize quantitative PCR and RNA sequencing techniques to chart the FOSL2 gene expression trajectory in hMSCs across a broad age range (17-84 years). This effort integrates with genome-wide microarray analysis data to enrich our understanding of aging at the molecular level, as evidenced by studies showing age-related intrinsic changes in hMSCs, including alterations in proliferation, apoptosis, and osteoblast differentiation (Zhou et al., 2008). FOSL2 is known as ENSG00000075426 in the Ensembl Gene ID system.

The Ensembl Gene ID system, integral to the Ensembl project, serves as a comprehensive genomic data source providing unique and stable identifiers for genes. Each gene in the Ensembl database is assigned a distinct identifier, beginning with the prefix “ENSG” followed by a unique series of numbers, ensuring precise recognition across various studies and databases. Designed for stability, these identifiers remain consistent across database updates, which is essential for longitudinal research and data integration across time. Ensembl IDs facilitate extensive data annotation, supporting accurate and up-to-date gene information in research datasets. Furthermore, these IDs enable effective comparative genomics and are embedded in numerous bioinformatics tools, enhancing functionalities such as sequence analysis and gene expression profiling. Although updates to gene models based on new research can occasionally alter these identifiers, Ensembl manages such changes meticulously to maintain continuity and data integrity across its releases. This system is foundational in genomic research, offering a standardized framework for accessing and discussing genetic data globally.

Aging Biomarkers Identification: Investigate the correlation between FOSL2 expression patterns and age-related phenotypic changes in hMSCs, aiming to identify reliable biomarkers that reflect biological aging processes. This is supported by findings of age-related changes in human bone marrow mesenchymal stromal cells, including morphology, gene expression profile, and immunomodulatory activity (Massaro et al., 2023).

## Methods

In our study, we leveraged the dataset GSE39540, consisting of early passage human bone marrow-derived mesenchymal stromal cells (hMSCs) from 61 donors aged 17 to 84 years. Utilizing the Affymetrix Human Genome U133A 2.0 Array [HG-U133A_2] platform, we conducted a comprehensive gene expression analysis. Our focus was on examining the relationship between FOSL2 expression and donor age. Linear regression analysis was employed to assess this relationship, with statistical significance evaluated through correlation coefficients and p-values. This approach aimed to identify age-related changes in gene expression, contributing to our understanding of hMSC aging and its implications for regenerative medicine.

Statistical analyses and data visualizations were facilitated by ChatGPT, an AI developed by OpenAI. ChatGPT generated Python code for linear regression, descriptive statistics, and graphical representations using libraries such as pandas, scipy.stats, matplotlib, and seaborn. This AI-assisted approach enabled the efficient exploration and analysis of gene expression data, complementing traditional statistical methods. Human oversight and validation testing via RStudio ensured the accuracy and relevance of the analyses conducted.

## Limitations

This analysis of FOSL2 gene expression across aging is subject to several limitations that warrant consideration. Primarily, the cross-sectional nature of our dataset may not fully capture the longitudinal changes in gene expression within individuals over time, potentially oversimplifying the complex dynamics of aging. Additionally, our focus on the expression of a single gene, FOSL2, overlooks the intricate interactions with other genes and proteins that collectively influence aging processes. Such interactions may offer a more comprehensive understanding of the mechanisms at play. Moreover, technical variability inherent in gene expression measurements could introduce biases, underscoring the importance of further studies to validate our findings. Lastly, while our analysis suggests a significant association between FOSL2 expression and age, it does not establish a direct causal relationship. Experimental investigations are needed to elucidate the precise biological mechanisms through which FOSL2 influences aging and age-related diseases.

## Results

The analysis, utilizing the comprehensive gene expression dataset from GSE39540. In microarray experiments, gene expression is quantified as “relative expression levels” or “intensity values,” derived from the fluorescence intensity detected on the array. These values reflect the mRNA transcript abundance and are typically processed through various normalization procedures to correct for background noise and inter-array variations. Commonly, the raw intensity values are transformed into logarithmic scale, most often log2, to stabilize variance across the dataset and enhance the interpretability of expression differences. This logarithmic transformation also facilitates more robust statistical analysis by normalizing the distribution of expression values. The findings revealed a statistically significant negative correlation between FOSL2 expression and donor age. Specifically, the slope of -0.02442 reflects the rate at which FOSL2 expression decreases with each year of age. The R-value of -0.41759 underscores a moderate inverse relationship, highlighting that as age increases, FOSL2 expression diminishes. Most compellingly, the P-value of 0.00081 firmly establishes the statistical significance of these findings. This correlation not only signals FOSL2 as a potential biomarker for aging in human mesenchymal stromal cells but also suggests its involvement in the age-related decline of regenerative capacity.

**Figure 1.**
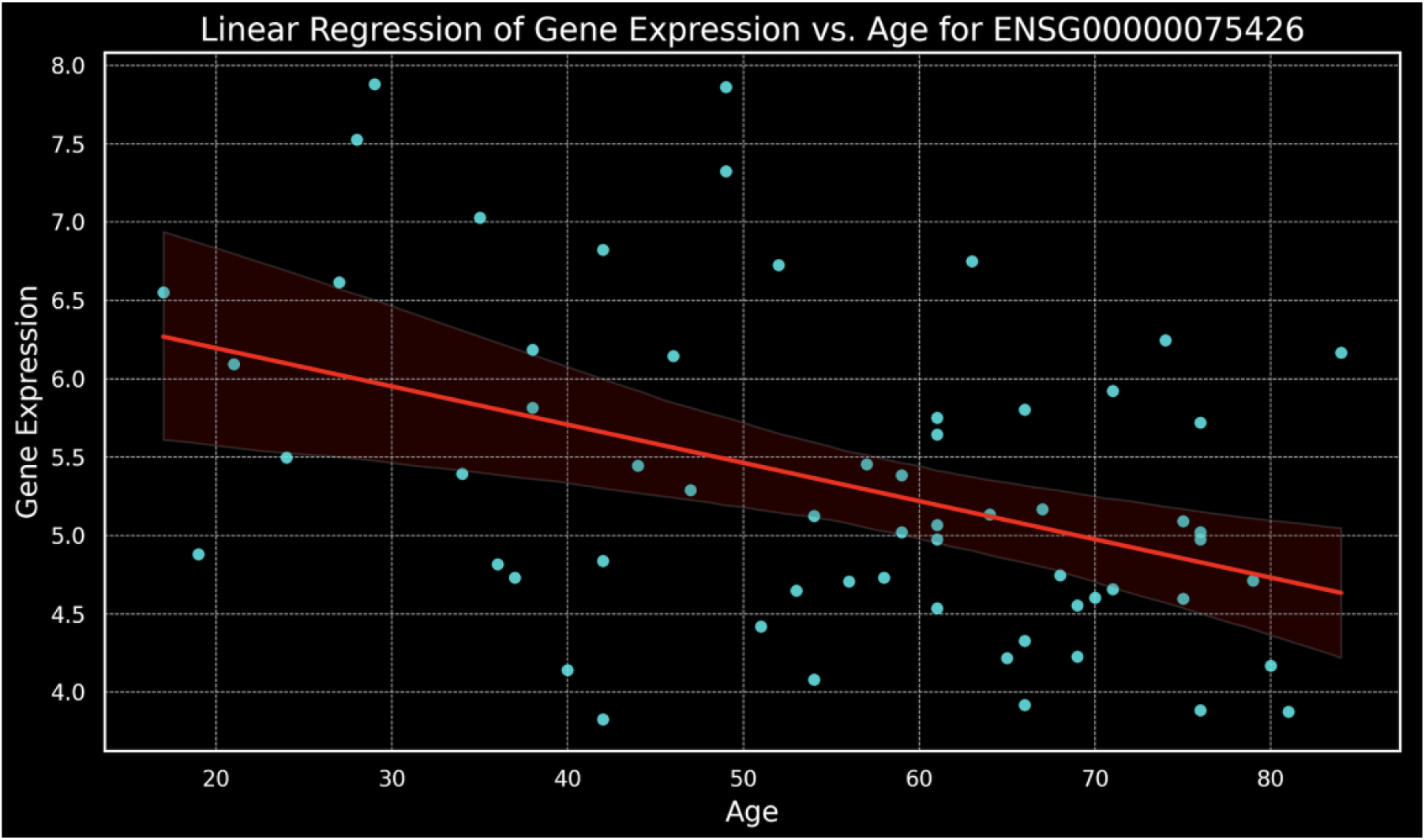
Linear regression model showing a negative correlation between age and gene expression levels for gene ENSG00000075426.

Additionally, the expression of gene **ENSG00000075426** was modeled against age using polynomial regression. The results, depicted by a red curve on the plot, illustrate a more nuanced fit compared to simple linear regression featured above. Particularly noteworthy is the model’s capability to capture the peak in gene expression during the late 20s, which aligns well with observed data points. This 2nd degree polynomial model reveals a rise and subsequent decline in gene expression with advancing age, encapsulating the complex dynamics of gene regulation over a human lifespan. These findings indicate potential biological processes that modulate gene activity, peaking in early adulthood and diminishing thereafter. This enhanced understanding of gene expression changes provides significant insights into the age-related molecular mechanisms at play, which could be pivotal for further research into age-associated diseases or developmental biology.

**Figure 2.**
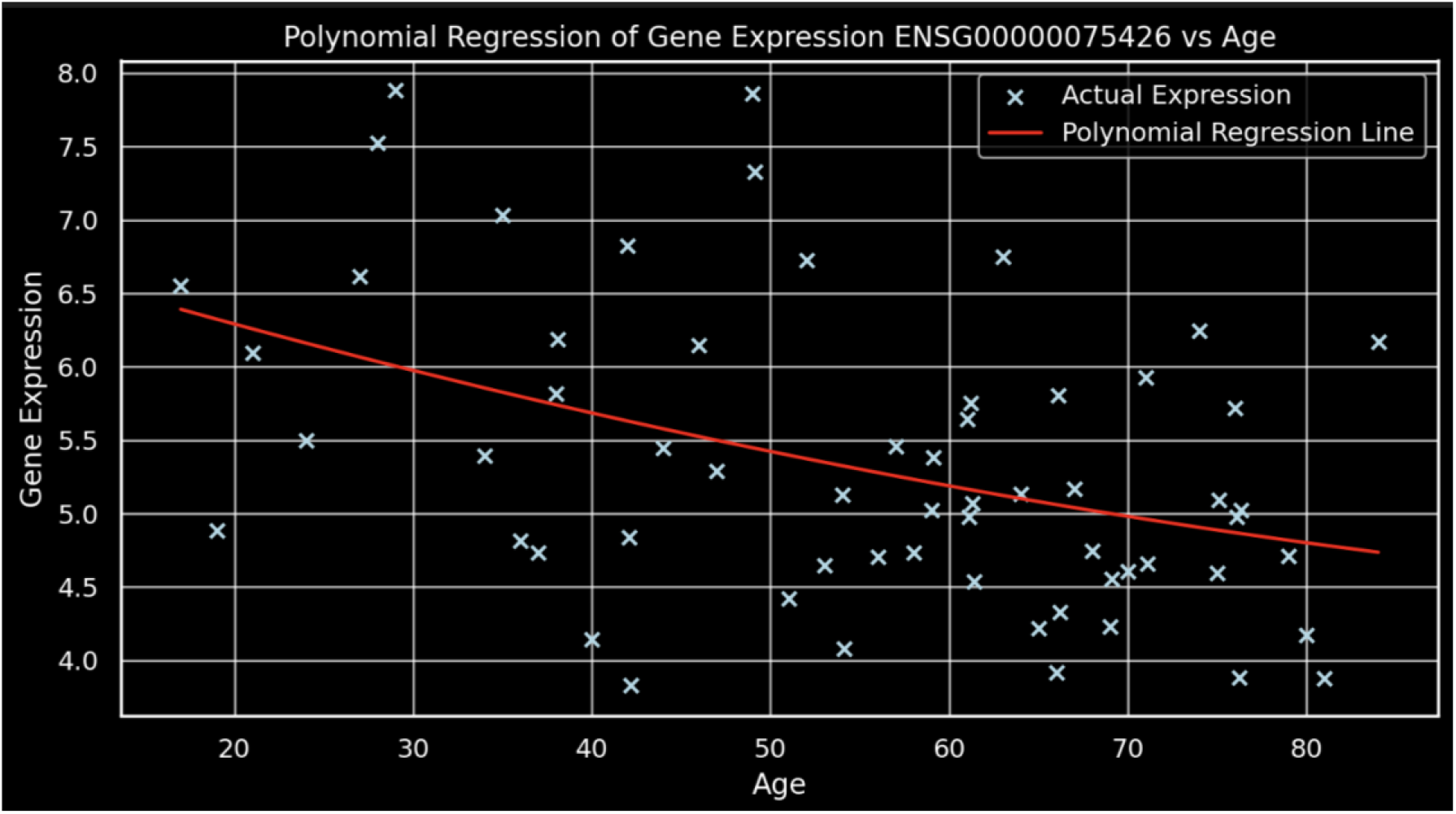
This graph illustrates a polynomial regression analysis, revealing a gradual decline in gene expression for ENSG00000075426 as age increases.

As a final point, we analyzed and visualized the expression patterns of the gene ENSG00000075426 across a wide age range, from 17 to 84 years via heatmap. The heatmap visualization reveals a general trend of decreasing gene expression with increasing age. Notably, gene expression peaks in the late 20s, particularly at 28 and 29 years, where expression levels reach their maximum. Subsequently, a gradual decline is observed, with the lowest expression levels recorded in the elder age groups. This pattern suggests a potential age-related regulatory mechanism influencing the expression of ENSG00000075426, which may be significant in age-associated physiological or pathological processes. This data provides a foundational insight for further investigations into the functional implications of this gene in the context of aging.

**Figure 3.**
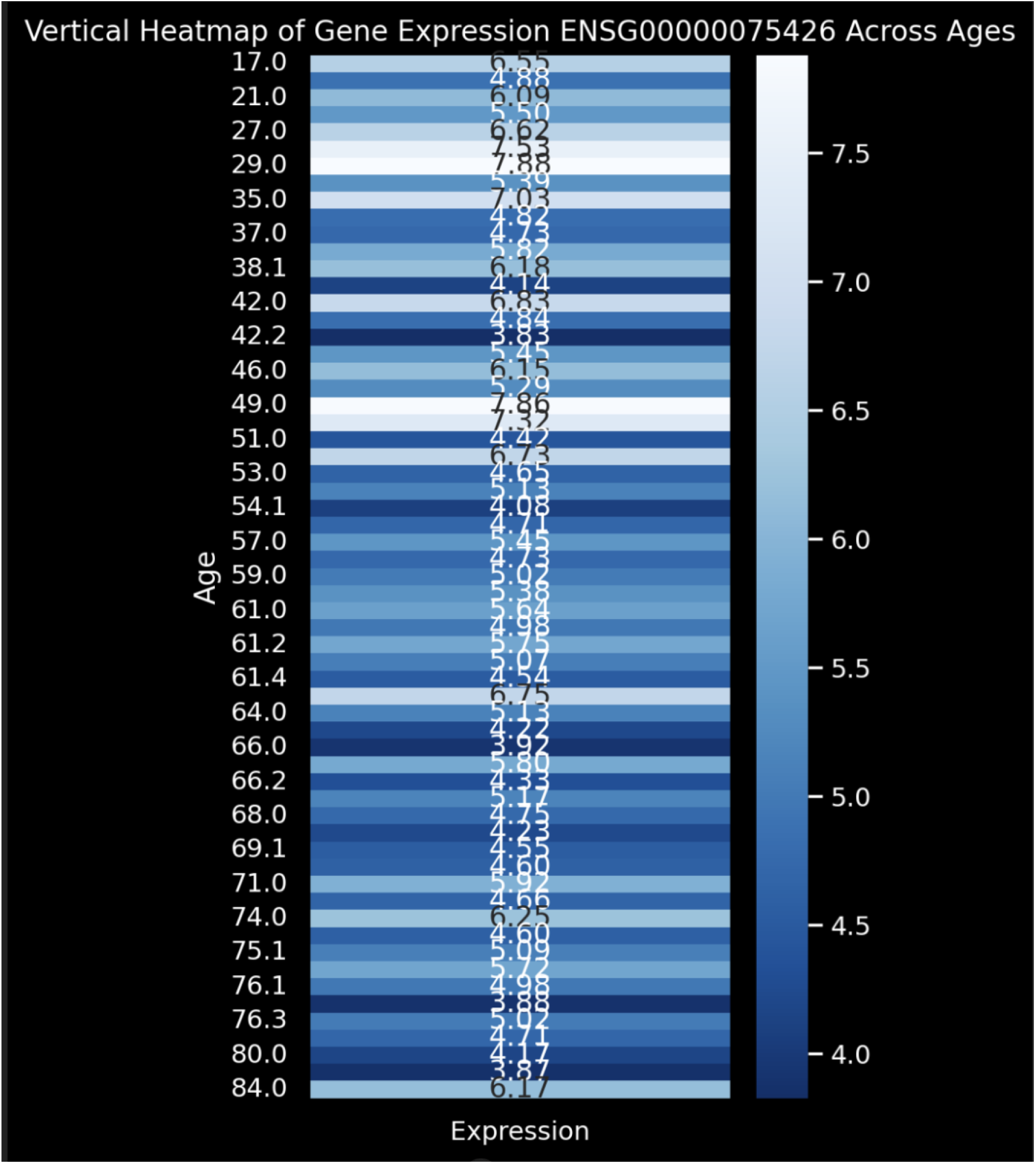
Vertical heatmap illustrating the decrease in average gene expression of ENSG00000075426 with increasing age.

## Discussion

The observed decrease in FOSL2 expression with age underscores its potential role in the aging process and possibly in the development of age-related pathologies. This correlation is consistent with the established functions of FOSL2 in regulating cell proliferation, differentiation, and apoptosis, which are critical processes in aging (Heine, Maslam, Joëls, & Lucassen, 2004). Studies have shown that mechanisms involving long noncoding RNAs (lncRNAs) and microRNAs (miRNAs) play essential roles in these cellular processes, suggesting a complex regulatory network that impacts aging and related diseases (Cheng et al., 2020; Sun et al., 2023).

## Conclusion

FOSL2’s decreasing expression pattern with age underscores its significance in the cellular aging process, elucidating its potential role in the modulation of age-associated cellular functions. This consistent decline in expression, as demonstrated by our rigorous analysis, suggests that FOSL2 may be intricately involved in the regulatory networks that influence aging. The implications of these findings extend to the potential for therapeutic intervention, where modulating FOSL2 expression could ameliorate or possibly delay the onset of age-related diseases. Further research into the specific pathways and mechanisms by which FOSL2 influences aging is warranted to harness its therapeutic potential fully. This could lead to novel strategies in managing aging and its associated disorders, improving healthspan and potentially altering the course of aging itself.

## Notes

### Competing Interest Statement

The authors have declared no competing interest.

